# Pharmacokinetics and Physiologically Based Pharmacokinetic Modeling of Mycobacteriophages: Insights into Pulmonary Distribution and Clearance

**DOI:** 10.64898/2026.01.27.702067

**Authors:** Rajnikant Sharma, Ramya Mahadevan, Sai Divyash, Shekhar Yeshwante, Saikumar Matcha, ChunFu Cheng, Hunter S. Talley, Alan A. Schmalstig, Pradeep Neupane, Sara E. Maloney Norcross, Anthony J. Hickey, Graham F Hatfull, Miriam Braunstein, Gauri G. Rao

## Abstract

Bacteriophage therapy is being explored as an alternate therapeutic approach for treating drug-resistant bacteria, including mycobacteria. However, rational phage dosing remains limited by scarce pharmacokinetic (PK) data and an incomplete understanding of tissue distribution. We performed dose-ranging studies in mice of three therapeutic mycobacteriophages (BPsΔ, ZoeJΔ, Muddy) after intravenous (IV) and intratracheal (IT) administration. All phages behaved similarly. IV dosing produced biphasic kinetics with non-proportional exposure and declining tissue-to-plasma ratios, indicating saturable uptake and elimination. IT delivery yielded monophasic profiles with ∼390-fold higher lung exposure and ∼490-fold lower plasma exposure, supporting inhaled therapy for pulmonary mycobacterial infections. Using BPsΔ data, we developed a mechanistic PBPK model incorporating transcytosis, saturable host clearance, plasma elimination, and lymphatic transport. The model accurately predicted ZoeJΔ and Muddy PK, enabled cross-species extrapolation, and showed that phage morphology influences disposition. This framework advances phage therapy toward model-informed, exposure-guided dose and route selection for multidrug-resistant bacterial infections.

## Introduction

The global rise in antimicrobial resistant (AMR) infections poses a serious threat to human health, with projections estimating over 39 million deaths from AMR bacterial infections by 2050^1,2^. Among the list of concerning AMR bacteria are nontuberculous mycobacteria (NTM), which are increasingly recognized as causes of pulmonary, skin, soft tissue, and disseminated infections^3^. Individuals with underlying lung conditions such as cystic fibrosis (CF), chronic obstructive pulmonary disease (COPD), and bronchiectasis as well as immunocompromised populations are at a particularly high risk of AMR NTM disease^4–12^. In the United States alone, over 59,700 new cases of NTM disease were reported from 2010 to 2019^13^.

*Mycobacterium abscessus* (Mabs), one of the most drug-resistant NTM species, remains exceptionally difficult to treat due to the lack of clinically validated treatment regimens, which makes it a critical target for new therapeutic development^14–19^. Bacteriophages (phages), which are viruses that infect and replicate within their specific bacterial hosts, are a promising alternative to antibiotics. Although phage therapy was first used in the early 20^th^ century, it was largely abandoned by Western medicine following the advent of antibiotics. However, the growing crisis of AMR has renewed interest in phage-based treatments, with recent compassionate-use cases demonstrating the potential for phage treatment being an alternative therapy. There are recent clinical cases of phages being used to treat AMR Mabs disease with favorable outcomes in 11 of 20 cases ^20,21^.

A limitation facing broader clinical application of phage therapy is the minimal understanding of key pharmacological principles for phage including: phage pharmacokinetics (PK), optimal phage delivery routes, and dosing strategies^22–25^. Unlike traditional small molecule drugs, phages are relatively large (∼100 nm), have a proteinaceous structure, and possess a unique “self-replicating” nature that distinguishes their PK. Most therapeutically used phages are double stranded DNA (dsDNA) tailed phages, with either contractile tails (myophages), long non-contractile tails (siphophages), or short stubby tails (podophages). These structural differences are likely to influence their PK.

In this study, our first objective was to characterize and compare the PK of three lytic mycobacteriophages (BPsΔ33HTH-HRM10 (BPsΔ), ZoeJΔ45 (ZoeJΔ), and Muddy), all of which are siphophages, in murine preclinical models. We evaluated their PK across different dose levels, routes of delivery (intravenous and intratracheal), and mouse strains. These phages were previously employed in compassionate-use cases to treat Mabs pulmonary disease with intravenous or inhaled (i.e., nebulization) delivery routes^20,26–29^. Our second objective was to develop a physiologically based PK (PBPK) model to gain mechanistic insights into phage biodistribution and elimination in plasma and key tissues (lung, liver, and spleen). The third objective was to evaluate the scalability potential of the PBPK model to predict PK of different phage families across different species (i.e., mouse, rat, monkey, and human).

## Results

We initiated PK analysis with BPsΔ, a mycobacteriophage used previously in multiple compassionate-use cases for treatment of pulmonary Mabs infections^20,26,28^. In most of these cases, BPsΔ was administered intravenously twice daily at ∼10^9^ PFU per dose. To define key determinants of phage disposition, we performed a stepwise evaluation of BPsΔ across varying dose levels, treatment conditions, and mouse strains. Additional experiments compared BPsΔ with ZoeJΔ and Muddy, while PBPK modeling integrated these findings into a predictive translational framework to enable interspecies scaling (**Fig.1**).

**Figure 1.**
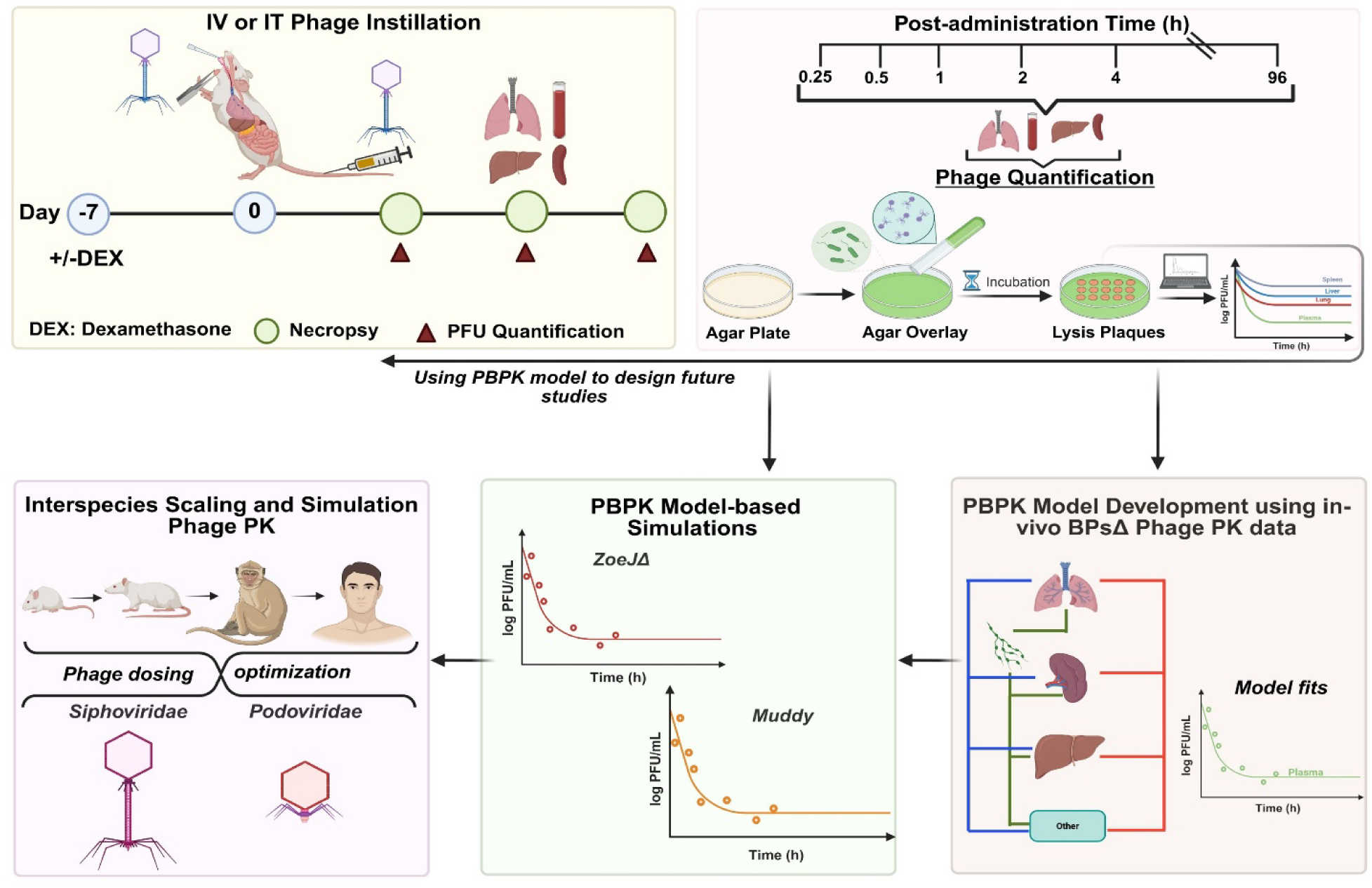
Visual Abstract. Overview of *in vivo* PK studies conducted in mice to assess phage distribution following intravenous (IV) or intratracheal (IT) administration in both dexamethasone treated and untreated (naïve) mice.

### Dose-Ranging PK and Tissue Distribution of IV Administered BPsΔ in DEX-Treated C3HeB/FeJ Mice

To assess BPsΔ PK and dose proportionality in vivo, we conducted a dose-ranging study following IV administration in uninfected, dexamethasone (DEX)-treated C3HeB/FeJ mice. This DEX-treatment model and mouse strain were selected as it is an established model for studying *M. abscessus* infection and drug treatment^30,31^. Three dose levels were evaluated, including high (4 × 10^12^ PFU/kg), medium (4 × 10^10^ PFU/kg), and low (4 × 10^7^ PFU/kg), with each administered as a single dose (**Fig. 2A**). Following IV administration, BPsΔ displayed a biphasic PK profile in plasma and all organs, characterized by an initial rapid distribution phase (up to 8 h) followed by a slower, prolonged elimination phase. For medium and high doses, phage titers remained at ≥ 2 log_10_ PFU/mL at 96 hours across all organs, whereas in the low-dose group, phages were detectable only up to 48 hours (**Fig. 2A**).

**Fig 2.**
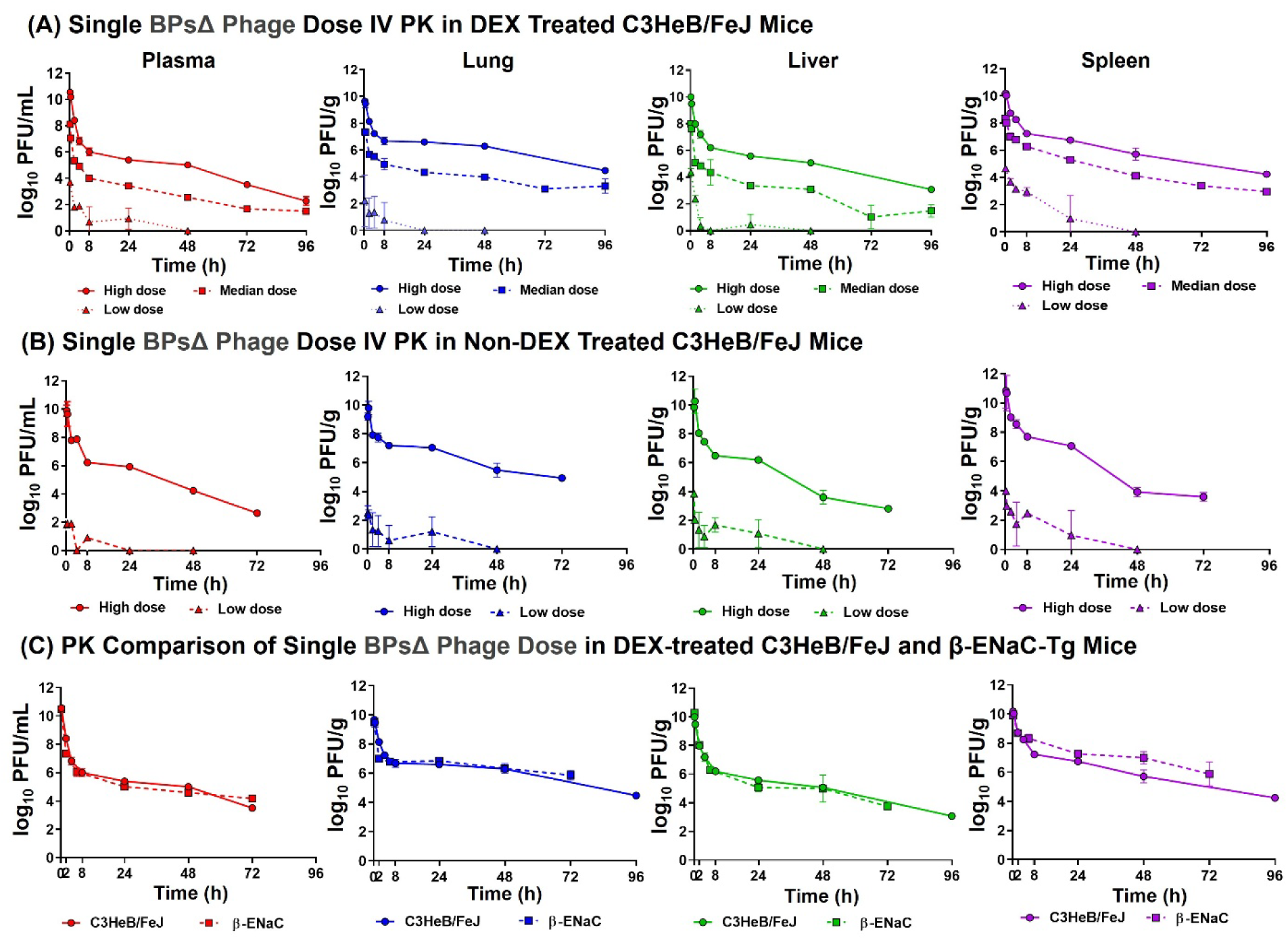
In vivo pharmacokinetics of BPsΔ phage following intravenous (IV) administration in dexamethasone (DEX)-treated and untreated mice. **Panel A–B:** Comparison of phage PK in DEX-treated (A) and non-DEX-treated (B) C3HeB/FeJ mice following single IV dose of BPsΔ at high (4 × 10¹² PFU/kg), medium (4 × 10¹⁰ PFU/kg), or low (4 × 10⁷ PFU/kg) concentrations. Phage titers (PFU) were quantified in plasma and organ homogenates (lung, liver, spleen, and lymph nodes) using the double agar overlay method. PK sampling was conducted over 96 hours for high and medium doses in DEX-treated mice, 72 hours in non-DEX-treated mice, and 48 hours for the low dose in both groups. Three mice were euthanized at each time point. **Panel C:** Single-dose IV PK comparison of BPsΔ in DEX-treated C3HeB/FeJ and β-ENaC-Tg mice at high (4 × 10¹² PFU/kg) dose. Sampling and quantification were performed as described above.

Area under the concentration–time curve from time 0 to infinity (AUC_0–∞_) for plasma, lung, liver, and spleen based on non-compartmental analysis (NCA) is summarized in **Table 1**. A 1,000-fold increase in dose (from 4 × 10^7^ PFU/kg for low dose to 4 × 10^10^ PFU/kg for medium dose) resulted in a >30,000-fold increase in AUC_0–∞_. Similarly, a 100-fold increase (from 4 × 10^10^ PFU/kg for medium dose to 4 × 10^12^ PFU/kg for high dose) led to a 155-fold increase in AUC_0–∞_. Dose-normalized AUC_0–∞_ values rose from 0.01 to 0.34 as the dose increased from low to high, (**Table 1**), suggesting saturation of phage clearance mechanisms at the higher doses.

**Table 1:**
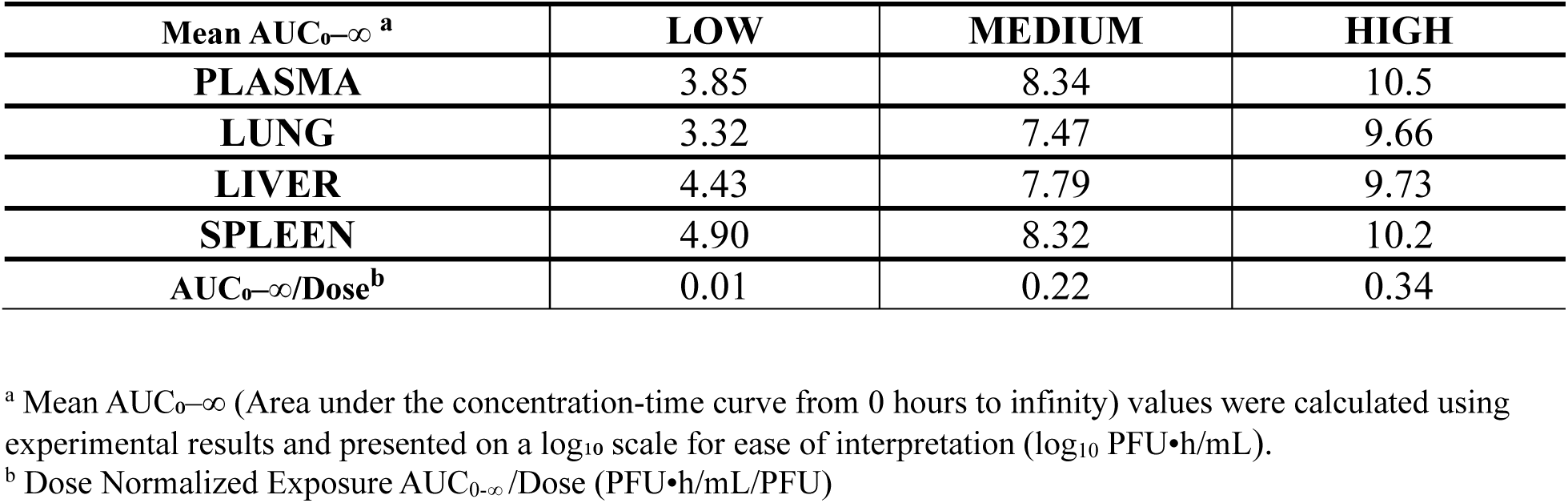
Comparison of Mean Exposure (AUC_0-∞_) and Dose Normalized Exposure (AUC_0-∞_ /Dose) Following Single Intravenous Administration of BPsΔ at High (4 × 10¹² PFU/kg), Medium (4 × 10¹⁰ PFU/kg), or Low (4 × 10⁷ PFU/kg) Dose

| Mean $AUC_{0-\infty}$ <sup>a</sup> | LOW | MEDIUM | HIGH |
| --- | --- | --- | --- |
| PLASMA | 3.85 | 8.34 | 10.5 |
| LUNG | 3.32 | 7.47 | 9.66 |
| LIVER | 4.43 | 7.79 | 9.73 |
| SPLEEN | 4.90 | 8.32 | 10.2 |
| $AUC_{0-\infty}/\text{Dose}$ <sup>b</sup> | 0.01 | 0.22 | 0.34 |

<sup>a</sup> Mean $AUC_{0-\infty}$ (Area under the concentration-time curve from 0 hours to infinity) values were calculated using experimental results and presented on a $\log_{10}$ scale for ease of interpretation ( $\log_{10}$ PFU•h/mL).
<sup>b</sup> Dose Normalized Exposure $AUC_{0-\infty}/\text{Dose}$ (PFU•h/mL/PFU)

Tissue-to-plasma AUC_0-∞_ ratios revealed marked differences in phage distribution across organs and doses. At the low dose, phage exposure was highest in the spleen (ratio = 11.1), followed by the liver (ratio = 3.74) and lung (ratio = 0.55), suggesting preferential uptake in spleen and liver. These ratios declined with increasing dose: at the medium dose (spleen: 0.96, liver: 0.29, lung: 0.14) and further at the high dose (spleen: 0.44, liver: 0.16, lung: 0.14), consistent with saturation of tissue uptake mechanisms.

### BPsΔ phage PK in DEX vs Non-DEX Treated C3HeB/FeJ Mice

We repeated the experiment using high and low IV doses in non-DEX treated C3HeB/FeJ mice to evaluate the impact of DEX treatment on phage PK. The resulting PK profile showed biphasic characteristics similar to that observed with the DEX treated group (**Fig. 2B**). Median plasma log_10_ AUC_0-∞_ values for the high dose were also similar between groups, DEX-treated (10.6 log₁₀ PFU·h/mL) and non-DEX-treated (10.7 log₁₀ PFU·h/mL) mice. No significant differences in phage exposure were observed in plasma and tissues, indicating that DEX treatment did not significantly affect BPsΔ PK at the high dose (Mann–Whitney U test, p > 0.05 for all organs; **Table 2**). Visual comparison of the low-dose PK profiles also suggests similar behavior between the two groups. However, statistical analysis was not possible as some of the PFU values were below the limit of quantification, resulting in a reduced sample size and unreliable statistical inference.

**Table 2:** Comparison of Phage Exposure (AUC_0-∞_) of BPsΔ Phage in Plasma, Lung, Liver and Spleen Following A Single Intravenous Dose in DEX and Non-DEX Treated C3HeB/FeJ mice & DEX Treated C3HeB/FeJ and β-ENaC-Tg Mice.

| Mean $AUC_{0-\infty}$<sup>a,b</sup> | C3HeB/FeJ | | β-ENaC-Tg |
| --- | --- | --- | --- |
|  | DEX Treated | Non-DEX Treated | DEX Treated |
| **PLASMA** | 10.6 (10.5 – 10.9) | 10.7 (9.60 – 10.8) | 10.7 (10.4 – 10.7) |
| **LUNG** | 9.81 (9.50 – 9.89) | 9.61 (9.59 – 10.3) | 9.60 (9.43 – 9.62) |
| **LIVER** | 9.72 (9.71 – 10.4) | 10.1 (9.70 – 11.1) | 10.3 (10.2 – 10.4) |
| **SPLEEN** | 10.3 (10.1 – 10.4) | 11.6 (10.2 – 12.0) | 10.2 (9.94 – 10.2) |

<sup>a</sup> Values are presented as median (range). $AUC_{0-\infty}$ (Area under the concentration-time curve from 0 hours to infinity) values were calculated using experimental results and presented on a $\log_{10}$ scale for ease of interpretation ( $\log_{10}$ PFU•h/mL).
<sup>b</sup> No significant difference in phage exposure was observed between DEX- and Non-DEX-treated groups across all organs (two-tailed Mann–Whitney U test, $p > 0.1$ ). No significant difference in phage exposure was observed between DEX treated C3HeB/FeJ and β-ENaC-Tg mice groups across all organs (two-tailed Mann–Whitney U test, $p > 0.1$ ).

### BPsΔ phage PK in DEX-Treated β-ENaC-Tg compared to C3HeB/FeJ mice

We repeated the high-dose IV BPsΔ study in uninfected, DEX-treated βENaC-Tg (Sccn1b-Tg) mice to compare the PK observed in C3HeB/FeJ mice with that observed in another mouse strain. βENaC-Tg mice exhibit hallmarks of CF lung disease, including mucus obstruction, hypersecretion, impaired mucociliary clearance, airway neutrophil inflammation, and reduced bacterial clearance^32,33^. Similar to DEX-treated C3HeB/FeJ mice, the PK profile in DEX-treated βENaC-Tg mice was biphasic, characterized by an initial rapid distribution phase followed by prolonged elimination. Phage exposure in plasma after high-dose administration was comparable to C3HeB/FeJ (10.6 log₁₀ PFU·h/mL) and β-ENaC-Tg (10.7 log₁₀ PFU·h/mL). No significant differences were observed in phage exposure in plasma or tissue between the two mouse strains (Mann–Whitney U test, p > 0.1; **Table 2**, **Fig. 2C**).

### Enhanced Lung Exposure via IT Administration

Given the clinical relevance of Mabs pulmonary disease, we next evaluated BPsΔ PK following IT administration in uninfected, DEX-treated C3HeB/FeJ mice at high and low doses. Although most mycobacteriophage treatments in clinical cases are currently administered intravenously, there are cases of BPsΔ being delivered via inhalation by nebulization^20^. IT administration showed a monophasic decline in the lung, liver, and spleen, which differs from the biphasic profile seen with IV administration in plasma and all organs (**Fig. 3**). Lung exposure was significantly higher with IT delivery; at the high dose, median lung AUC₀–∞ increased from 9.81 log_10_ PFU·h/mL (IV) to 12.4 log_10_ PFU·h/mL (IT). In contrast, plasma exposure was reduced (IV vs. IT: 10.6 log_10_ PFU·h/mL vs. 7.97 log_10_ PFU·h/mL). Liver and spleen exposures were similar between IV and IT routes (**Supplemental Table 1**, **Fig. 3A & B**). At the low dose, IT administration also resulted in higher lung concentrations compared to low-dose IV delivery. Plasma concentrations following low-dose IT administration were below the limit of quantification across all time points, reflecting minimal systemic exposure (**Fig. 3B**).

**Fig 3.**
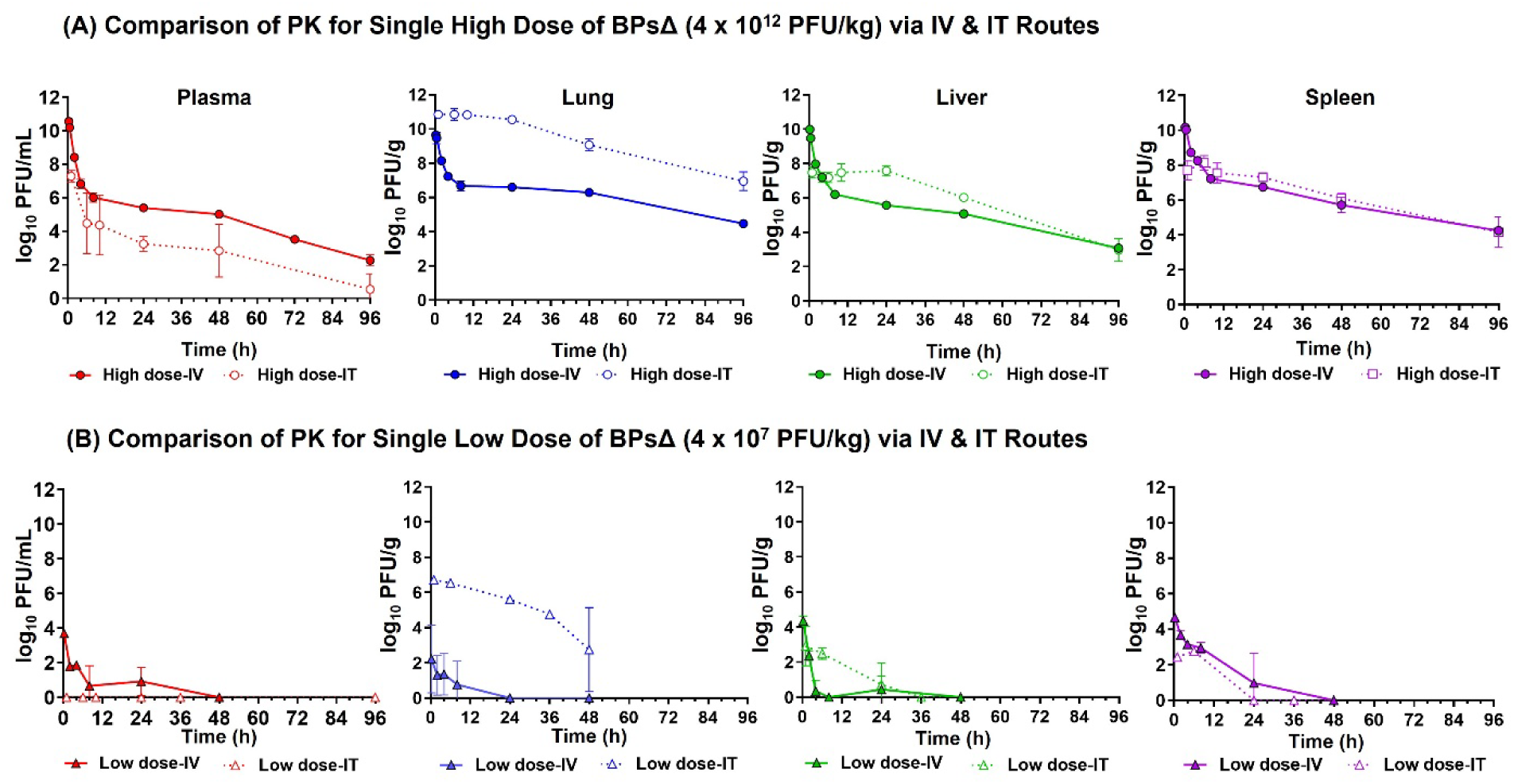
Comparison of *in vivo* pharmacokinetics following administration of single dose of BPsΔ phage via IV and IT routes in DEX-treated C3HeB/FeJ mice. Mice received a single high (4 × 10¹² PFU/kg) (**A**) or low (4 × 10⁷ PFU/kg) (**B**) dose of BPsΔ via intravenous (IV) or intratracheal (IT) routes. Phage titers were quantified in plasma and organ homogenates (lung, liver, and spleen) using the double agar overlay method. PK sampling was conducted over 96 hours for the high dose and 48 hours for the low dose. Three mice were euthanized at each time point.

### Comparison of PK of BPsΔ, ZoeJΔ, and Muddy Phages

To compare BPsΔ PK with other clinically relevant phages, we repeated high-dose experiments with ZoeJΔ and Muddy mycobacteriophages. Following IV administration, all three phages showed comparable exposure in plasma, lung, liver, and spleen, with no statistically significant differences. Across phages, splenic exposure was highest, followed by exposure achieved in the liver. Muddy and ZoeJΔ exhibited splenic AUC values that exceeded their plasma exposure after IV administration, (**Supplementary Table 1, Fig. 4A**).

**Fig 4.**
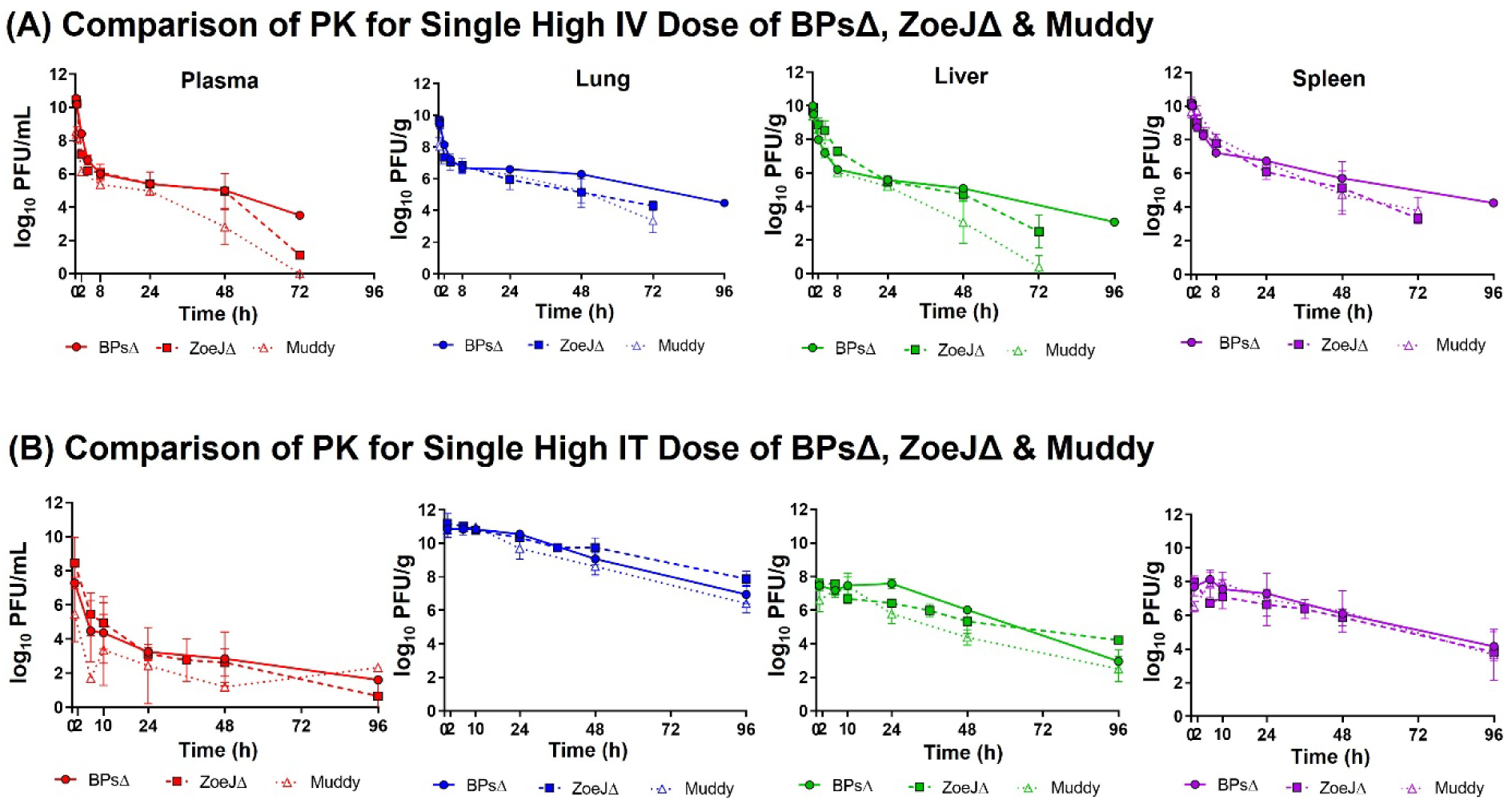
Comparison of PK following administration of single high dose (4 × 10¹² PFU/kg) of BPsΔ, ZoeJ Δ, and Muddy phage via IV or IT routes in DEX-treated C3HeB/FeJ mice. **(A)** In vivo PK following a single high IV dose of BPsΔ and ZoeJ Δ and Muddy. **(B)** In vivo PK following a single high IT dose of BPsΔ and ZoeJ Δ and Muddy. Phage titers were quantified in plasma and organ homogenates (lung, liver, and spleen) using the double agar overlay method. Sampling was conducted over 96 hours, with three mice euthanized at each time point.

After IT administration, no significant differences in exposure were observed among the three phages in plasma, lung, liver, or spleen. IT delivery consistently resulted in elevated lung exposure compared to IV administration (high-dose AUC_0–∞_ IT vs. IV: BPsΔ: 12.4 log_10_ PFU·h/mL vs. 9.81 log_10_ PFU·h/mL; ZoeJΔ: 12.4 log_10_ PFU·h/mL vs. 9.90 log_10_ PFU·h/mL; and Muddy: 12.3 log_10_ PFU·h/mL vs. 8.52 log_10_ PFU·h/mL). After IT delivery, among organs, lung exposure was greatest, followed by spleen and liver, with this trend consistent across all three phages (**Supplementary Table 1, Fig. 4B**).

### PBPK Modeling of BPsΔ Phage

To characterize BPsΔ disposition, we developed a PBPK model using a naïve pooled approach with phage concentration-time data from low, medium, and high doses administered via IV or IT routes. The model included compartments for organs of interest with regards to phage disposition (plasma, lung, liver, spleen, and lymph node **Fig. 5A**). Tissue compartments were subdivided into vascular, interstitial, and cellular spaces and linked through plasma and lymph flow (**Fig. 5B, C)**.

**Fig 5.**
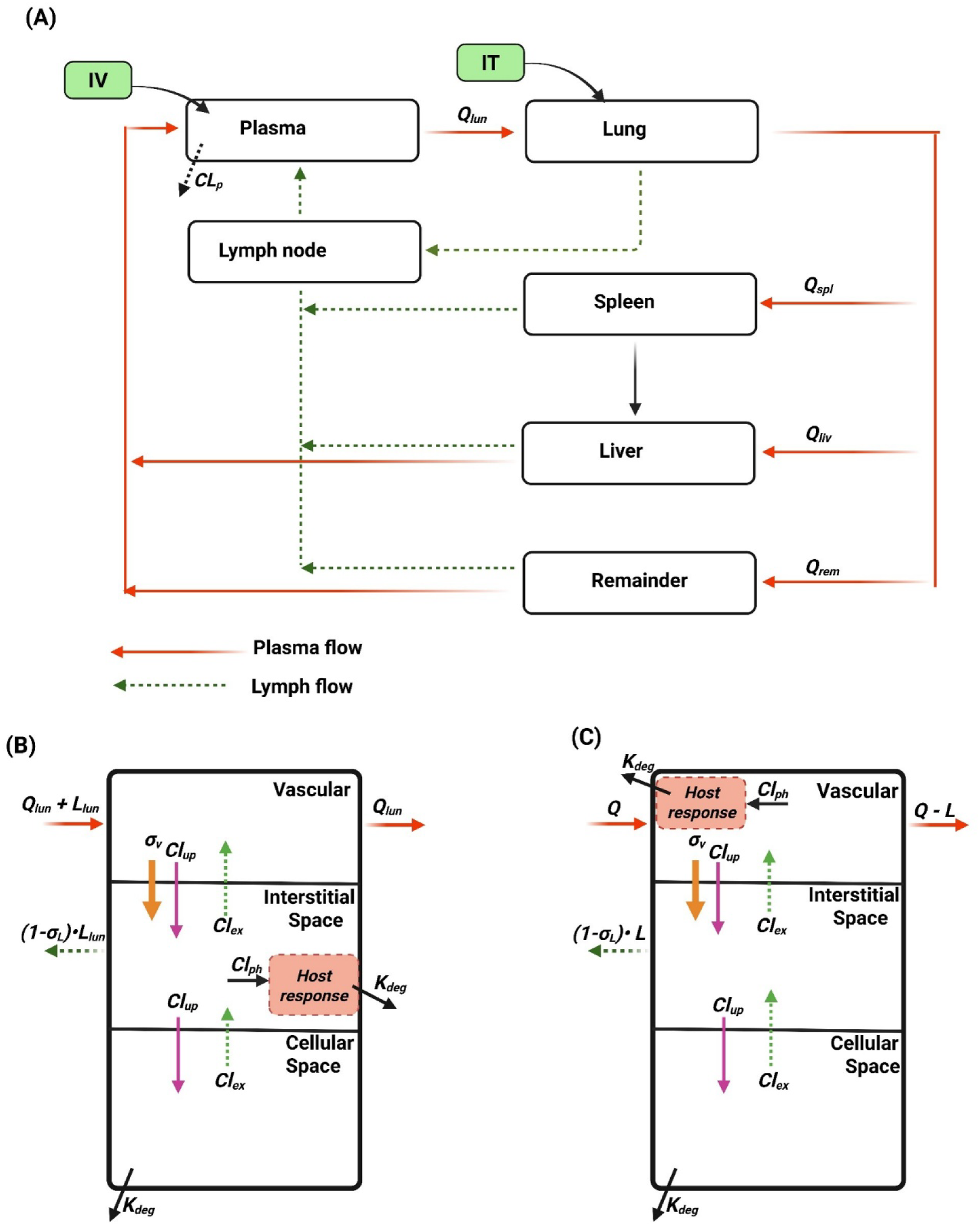
PBPK Model Schematic for Bacteriophages. **(A)** Overview of the physiologically based pharmacokinetic (PBPK) model, showing compartments for lung, liver, spleen, and remaining organs lumped as ‘Reminder’. Each tissue compartment is subdivided into vascular, interstitial, and cellular spaces. Bacteriophage transport occurs via blood circulation (solid arrows) and lymphatic flow (dashed arrows). *Q_org_* and *L_org_* represent organ plasma flow and lymph flow respectively; *CL_p_ denotes* plasma clearance. (**B-C**) Tissue-level PBPK structures for lung (**B**), liver, and spleen (**C**) including vascular, interstitial and cellular spaces. Phages undergo endocytosis (*Cl_up_*) from vascular to cellular space and exocytosis (*Cl_ex_*) from cellular to vascular space. Convective transport (*σ_v_*) moves phages from vascular to interstitial space, while lymphatic drainage (*σ_L_*) transfers them from interstitial space to lymph nodes. Host response includes phagocytic uptake (*Cl_ph_*) by macrophages in lung interstitial space and in vascular spaces of liver and spleen. Tissue-level degradation (*K_deg_*) occurs due to host response and lysosomal activity within cellular spaces.

Phages exiting interstitial spaces via lymphatic drainage were routed to the lymph node before returning to plasma. Supporting a possible role of lymphatics in phage distribution, we quantified phage levels in lymph nodes after IV or IT administration and observed substantial phage exposure with a biphasic profile after IV and a monophasic profile after IT (**Supplementary Fig 1**). Physiological parameters are listed in **Supplementary Table 2**, and governing equations are provided in **Supplementary Materials**.

Phage transport between vascular and interstitial spaces can occur through two distinct mechanisms. The first mechanism is convective transport via paracellular pores, based on the two-pore hypothesis, assuming vascular endothelium porosity with small and large pores^34^. Since the maximum small pore radius is 9 nm and phages exceed this size, phage transport would occur through large pores^34^. The model predicted convective transport was governed by a vascular reflection coefficient (σ_vas_ = 0.99), representing the fraction of phages in the tissue vascular space unable to cross via the pores. Convective transport could also involve interstitial fluid drainage into the lymphatic system and then to vascular space, the model included a lymphatic reflection coefficient (*σ*_*lym*_ = 0.2), indicating partial resistance due to physical size and electrostatic interactions with the extracellular fluid matrix. The second transport mechanism for phages to move between vascular and interstitial spaces is transcytosis across the vascular endothelium, which is supported by phage uptake studies performed in cultured human cells^35–37^. This mechanism would involve endocytosis (*Cl*_*up*_) and exocytosis (*Cl*_*ex*_). Model estimates predicted phage uptake by cells occurred at 2.37 mL/h (*Cl*_*up*_), while exocytosis (*Cl*_*ex*_) varied by tissue, being higher in spleen and liver (4.99 mL/h) and lower in lung (1.92 mL/h). From the interstitial space, phages could also be internalized by cells via endocytosis and degraded by cellular lysosomes (*K*_*degr*_).

A central feature of the model was incorporating host response–mediated saturable phage uptake. This mechanism likely drives the dose-dependent clearance, as phage degradation occurs after uptake. This saturable process explains the non-proportional exposure and the shift in tissue-to-plasma AUC_0–∞_ ratios as the dose increases. In the lungs (**Fig. 5B**), uptake was modeled in the interstitial sub-compartment to reflect macrophage-mediated endocytosis and degradation. In the liver and spleen (**Fig. 5C**), uptake was modeled in the vascular sub-compartment for macrophages such as Kupffer cells (in the liver) and splenic macrophages contributing to phage uptake and clearance. This process was described by:

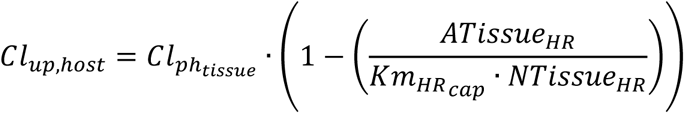

where 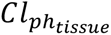 (h^−1^) is the maximum uptake rate constant in the tissue, *ATissue*_*HR*_is the phage amount in host cells, 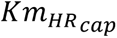 (PFU/cell) is uptake capacity per host cell, *and NTissue*_*HR*_(cells) is the number of host cells (i.e., number of macrophages).

Estimates for tissue uptake indicated spleen was the fastest (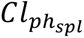 38.0 h⁻¹), followed by liver (5.17 h⁻¹) and lung (3.0 h⁻¹). It is noteworthy that high spleen exposure was observed across IV and IT routes of administration **(Supplementary Table 1)**. Estimated phage uptake capacity per host cell 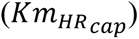 was 3.85 log_10_ PFU/cell.

Host-mediated phage degradation via immune cell clearance by cellular mediated lysosomal processing varied by route: IV dosing predicted faster degradation (*K*_*degr*_ =1.54 h⁻¹) compared to IT dosing (*K*_*degr*_ =0.12 h⁻¹). To accurately capture lung kinetics, a lung-specific host immune mediated degradation rate constant was incorporated. This rate remained low 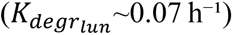 regardless of route, supporting prolonged lung exposure (**Fig. 4A & B and Fig. 6A & B**).

**Fig 6.**
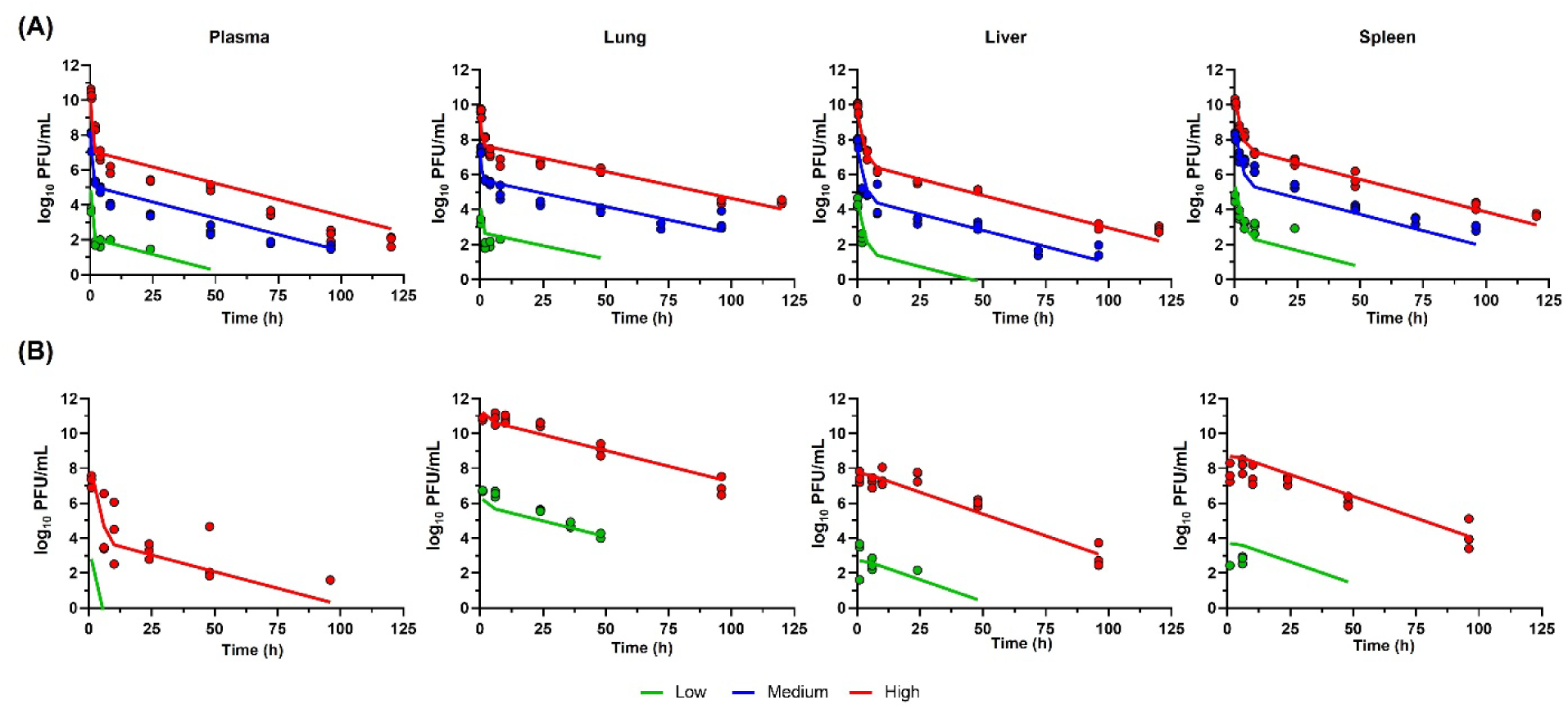
Simultaneously fitted concentration-time profiles of BPsΔ in plasma, lung, liver and spleen following IV (A) and IT (B) administration using the PBPK model after parameter optimization incorporating host response. For IV administration, three different dose levels were evaluated: high (4 × 10¹² PFU/kg), medium (4 × 10¹⁰ PFU/kg), and low (4 × 10⁷ PFU/kg). For IT administration high and low were used. Solid line represents the PBPK model predictions, while closed circles denote observed experimental data.

Systemic phage clearance that was not explained by tissue-specific host-mediated mechanisms was modeled as plasma clearance (*Cl_p_*). The model revealed substantial route-dependent differences; after IV administration, plasma clearance was 10.9 mL/h, whereas IT administration resulted in markedly higher plasma clearance (472 mL/h). Overall, the PBPK model accurately captured observed data for both IV and IT administration at different dose levels (**Fig. 6A & B**) with precise parameter estimates (**Supplementary Table 3**).

### Model-Based Simulations of ZoeJΔ and Muddy PK

We simulated the PK of ZoeJΔ and Muddy phages, which are morphologically similar to BPsΔ, using final parameter estimates from the PBPK model. The model successfully simulated plasma and tissue PK following both IV and IT administration, with majority of observed data falling within the 90% prediction interval (**Fig. 7 for ZoeJ**Δ**, Supplementary Fig. 2 for Muddy**). With the exception of high variability in plasma PK after IT administration, the model-based simulations captured the median profile.

**Fig 7.**
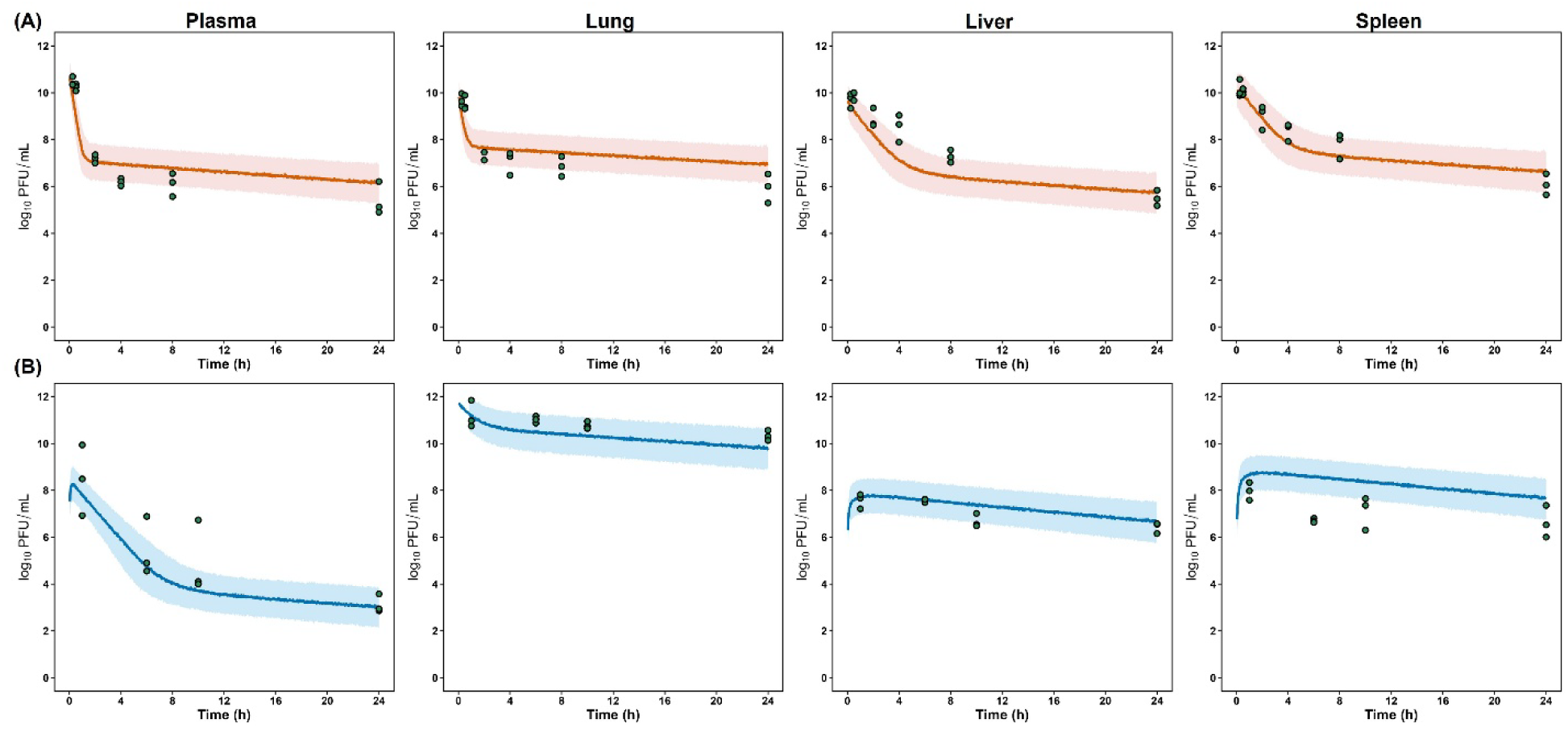
PBPK model-based simulations of pharmacokinetics following a single high-dose (4 × 10¹² PFU/kg) administration of ZoeJΔ phage via intravenous (IV) and intratracheal (IT) routes. Simulated concentration-time profiles in plasma, lung, liver, and spleen are shown for IV (**A**) and IT (**B**) administration. Solid lines represent the median predictions, with the shaded areas indicating the 5th to 95th percentile range. Symbols correspond to observed experimental data.

### Interspecies Scaling of the PBPK Model

We next evaluated the ability of our PBPK model to simulate the PK of other phages, including podophages because the model was developed using data from siphophages. Phage PK data from four published studies (three involving podophages^38–40^ and one with a siphophage^41^) were digitized (**Supplementary Table 4**)^38–41^. Using species-specific physiological parameters and allometric scaling of plasma clearance, the PBPK model developed in mice successfully captured PK across species, including uninfected mice (**Fig. 8**), rats (**Fig. 9 A, B & C**), and monkeys (**Fig. 9 D &E**). Notably, the PBPK model developed using siphophage data accurately predicted PK for podophages after parameter optimization to account for size- and morphology-dependent differences in this phage family.

**Fig 8.**
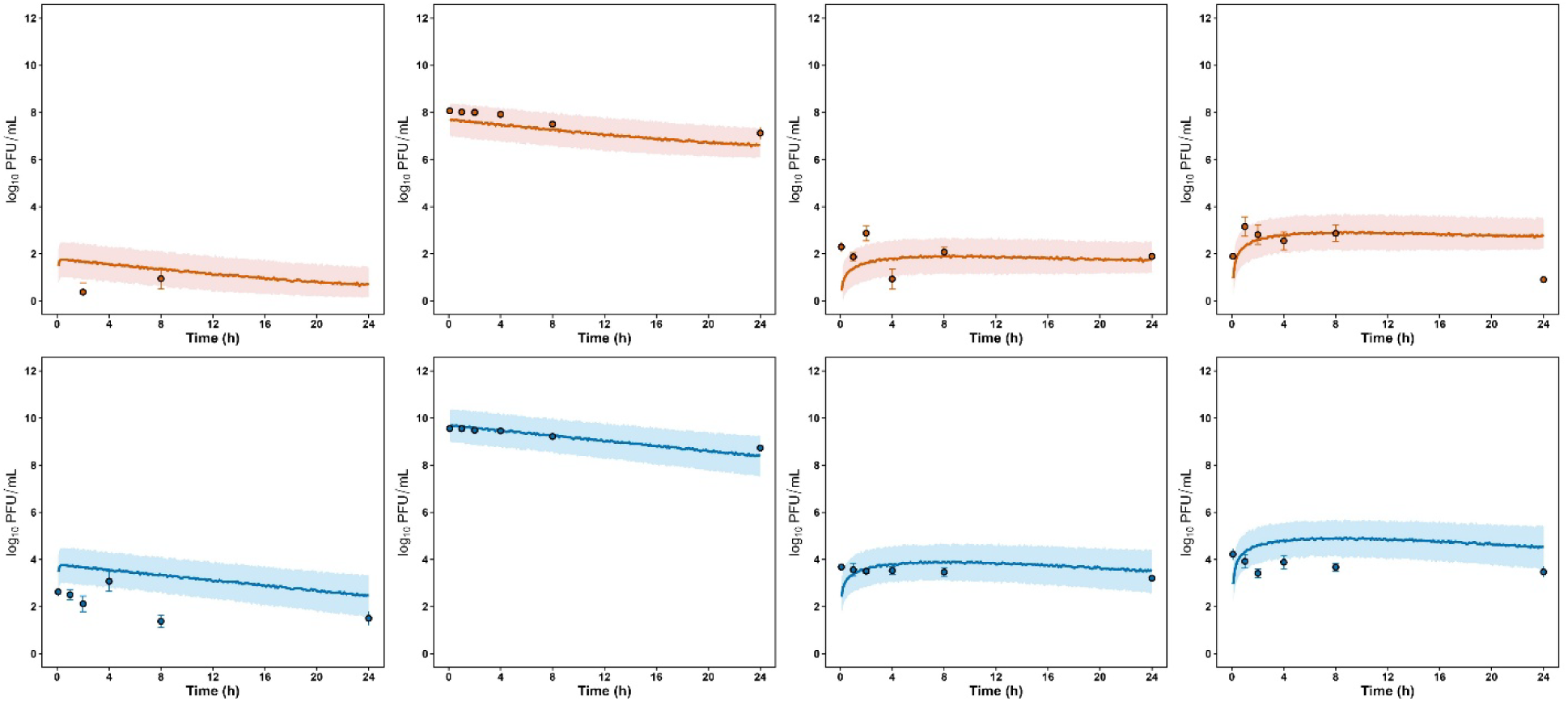
Model based simulations of single IT dose PK for *Pseudomonas aeruginosa phage, PEV31 for two different dose levels,* 10^7^ PFU (A) and 10^9^ PFU (B) in uninfected mice. The solid line represents the median and 5th–95th percentiles of predictions in shaded area. The symbols represent the observed data digitized from Chow et al 2021.

**Fig 9.**
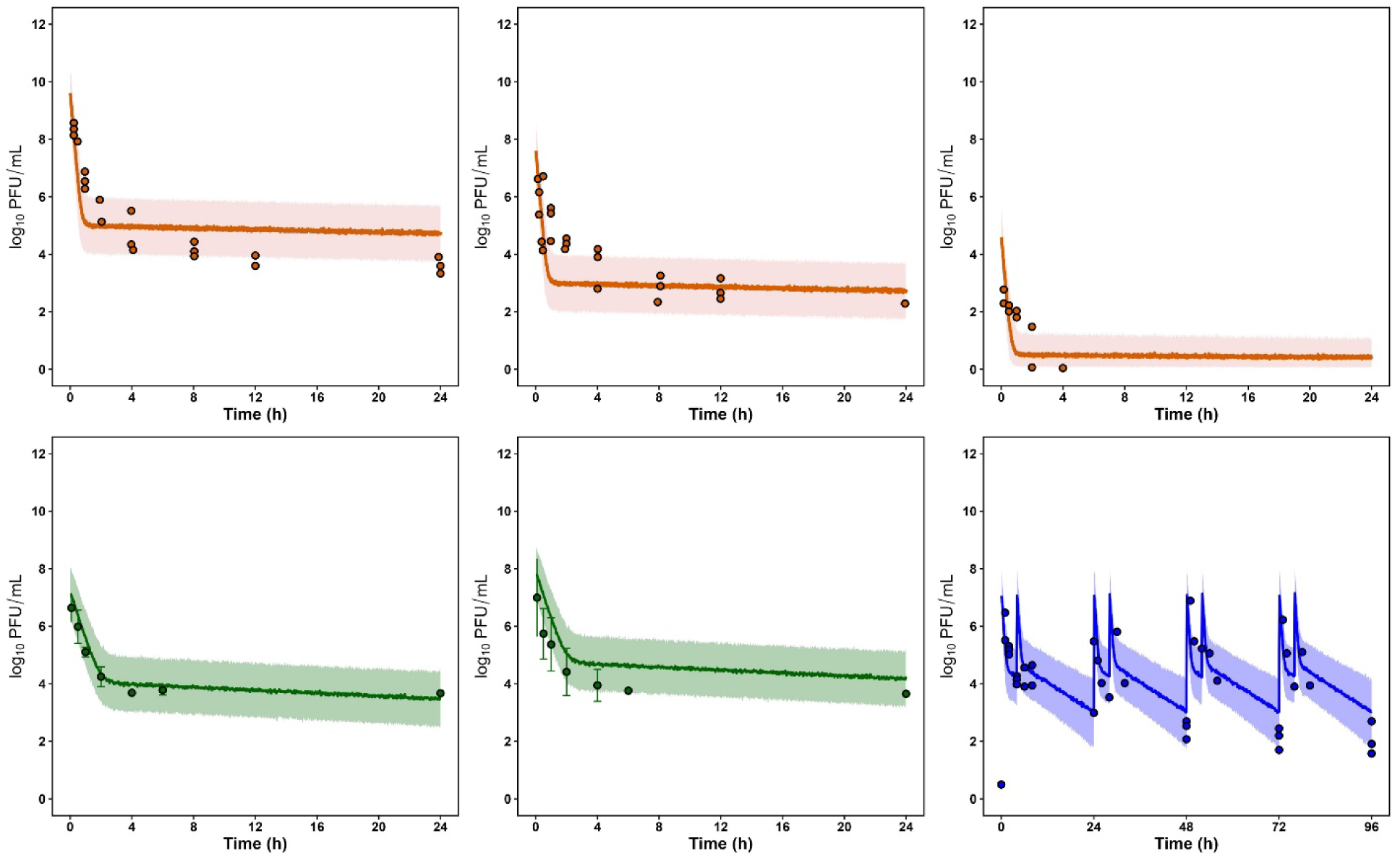
A, B & C: PBPK Model based simulations of single IV dose PK for *Pseudomonas aeruginosa phage, øPEV20 for three different dose levels,* 11 log10 PFU/rat. **(A), 9 log10 PFU/rat (B) and 6 log10 PFU/rat (C) in uninfected rats.** The solid line represents the median blood concentration and 5th–95th percentiles of predictions in shaded area. The symbols represent the observed data digitized from Lin et al 2020. **D & E: Model based simulations of single IV dose PK for *Salmonella phage, SE_SZW1 for two different dose levels,* 10**^9^ **PFU/kg (D) and *5*·10**^9^ **PFU/kg (E) in uninfected monkeys.** The solid line represents the median blood concentration and 5th–95th percentiles of predictions in shaded area. The symbols represent the observed data digitized from Tan et al 2023. (**F**) **Model based simulations of 1 × 10**^11^ **PFU IV bolus dose twice daily,4 h apart PK for phage cocktail*, LBP-EC01* in patients with uncomplicated UTI.** The solid line represents the median blood concentration and 5th–95^th^ percentiles of predictions in shaded area. The symbols represent the observed data digitized from Kim et al 2024.

It is reported that podophages, which have smaller short stubby tails compared to siphophages, are associated with higher cellular uptake (i.e, intracellular concentrations) and faster plasma clearance compared to siphophages^42,43^. Model parameters were optimized by adjusting exocytosis rates from cellular to vascular spaces and increasing plasma clearance (**Supplementary Table 4**) to account for these characteristics. With this optimization, the model successfully described plasma and tissue PK of PEV31 after IT administration of 10⁷ and 10⁹ PFU doses in mice (**Fig. 8**)^38^ and øPEV20 in rats at three IV dose levels (11, 9, and 6 log₁₀ PFU/rat) (**Fig. 9 A, B and C**)^39^.

In contrast, the *Salmonella* siphophage SE_SZW1 was well described in monkeys using only allometrically scaled plasma clearance, without additional parameter optimization, which likely reflects the fact that SE_SZW1 and BPsΔ, the latter being the phage used to develop the PBPK model, are both siphophages. Plasma PK at two dose levels (10⁹ and 5 × 10⁹ PFU/kg) matched model simulations (**Fig. 9 D & E**)^41^.

Finally, the PBPK model was scaled to humans using species-specific physiological parameters, and allometrically scaled plasma clearance to assess translational utility. Plasma PK data from humans treated for urinary tract *E. coli* infection with an *E. coli* phage cocktail (LBP- EC01) was used. The cocktail included T3 and T7 phages; therefore, podophage-like PK was assumed, and *podophage*-specific parameter optimization applied (**Supplementary Table 4**). The model accurately captured plasma concentration–time profiles in patients receiving 1 × 10¹¹ PFU IV twice daily, spaced 4 hours apart (**Fig. 9F**)^40^.

## Discussion

In a recent retrospective study of 100 difficult-to-treat bacterial infections, adjunctive phage therapy produced clinical improvement in 77.2% and bacterial eradication in 61.3% of cases^44^. In compassionate-use cases, mycobacteriophage therapy has similarly shown potential for treating multidrug-resistant NTMs, including Mabs, with favorable clinical and microbiological outcomes in 11 of 20 treated cases^20^. To date, only a few phages, most commonly BPsΔ, ZoeJΔ, and Muddy have been used therapeutically, administered intravenously, via aerosolization, or through both routes, typically at doses of ∼10⁹ PFU (∼ 2×10^7^ PFU/kg) twice daily^28^. Despite increasing clinical use of phages being reported, critical gaps persist in defining optimal dosing, route, frequency, and site-of-infection exposure. Pharmacokinetic data remain especially limited and are lacking altogether for BPsΔ, ZoeJΔ, and Muddy, leaving current IV and nebulized doses largely empirical. Quantitative PK studies are therefore essential to guide rational phage therapy design.

Here, we provide a comprehensive quantitative analysis of mycobacteriophage disposition showing: (i) nonlinear, saturable PK after IV dosing; (ii) biphasic tissue kinetics after IV administration versus monophasic profiles after IT dosing; (iii) enhanced lung exposure with IT dosing, yielding ∼390-fold higher lung exposure and ∼490-fold lower systemic exposure compared with IV administration; (iv) phage PK is unaffected by DEX treatment or the CF disease-like features of β-ENaC-Tg mice; (v) comparable PK characteristics across all siphophages within their respective IV and IT administration groups; and (vi) development of a mechanistic PBPK model incorporating endocytosis/exocytosis, saturable host-response clearance, and lymphatic flow that predicts PK across phage families and species, identifies size- and morphology-dependent PK, and informs rational dose and route selection for clinical translation.

Our results indicate that tissue uptake and phage elimination are both saturable processes. The IV dose-ranging study revealed a non-proportional increase in exposure with increasing phage doses indicating non-linear PK likely driven by saturation of clearance mechanisms. This aligns with previous *Pseudomonas* phage øPEV20 PK studies in rats^39^. Such non-linearity is consistent with phage clearance mediated by mononuclear phagocytic system (MPS), where macrophage-mediated phagocytosis in the liver, spleen, and lungs serves as a principal elimination pathway^45^. The decreasing tissue-to-plasma ratios at higher doses supports saturable uptake, consistent with concentration-dependent uptake previously reported for M13-RGD8 phage^46^.

Following IV administration, plasma, lung, spleen, and liver exhibited biphasic kinetics, with an initial rapid distribution phase followed by a prolonged elimination phase, as reported for IV delivery of øPEV20 phage in rats. Despite systemic dosing, phage portioning to the lung was substantial, with lung exposure (3.32 to 9.66 log_10_ PFU·h/mL) remaining close to plasma exposure (3.85 to 10.5 log_10_ PFU·h/mL) across all dose levels (**Table 1**). In contrast, IT administration yielded monophasic profiles across all three organs, as was previously observed with IT PEV31 delivery in mice^38^. Furthermore, IT delivered BPsΔ phage resulted in ∼390-fold higher lung exposure and ∼490-fold lower plasma exposure relative to IV dosing (**Fig. 3)**.

Importantly, our PK findings appear to apply to multiple mouse models. Because MPS functions in phage clearance and DEX immunosuppression is critical to maintain Mabs infection in the C3HeB/FeJ mouse model^30^, we evaluated DEX effects on phage PK^45^. No significant differences were observed between DEX-treated and untreated groups, consistent with reports that prednisone does not acutely alter MPS-mediated elimination of radiolabeled human IgG aggregates^47^. Similarly, phage exposure in DEX-treated β-ENaC-Tg mice, modeling CF-like airway pathology^32,33^, was comparable to C3HeB/FeJ mice, indicating that mucus hypersecretion and impaired mucociliary clearance in β-ENaC-Tg mice do not significantly alter phage PK in plasma and tissues after IV or IT dosing.

Most prior phage PK studies rely on non-compartmental analysis, which cannot resolve mechanisms of phage disposition. In contrast, PBPK models integrate systemic and tissue-level PK with biological mechanisms. PBPK models have been applied to characterize PK of large molecules, including nanoparticles, monoclonal antibodies, and AAVs^34,48,49^. Phages share multiple PK characteristics with nanoparticles, including MPS-mediated clearance, biphasic IV and monophasic IT profiles, lymph-mediated distribution, and size/morphology dependent differences in PK^34,50^. Notably, our study is the first to quantify phage concentrations in lymph, revealing a role in phage distribution. Importantly, the PBPK framework supports interspecies scaling to project human PK and guide dosing optimization^51^.

Compared to a recent phage PBPK model that assumed generalized permeability-limited distribution and MPS clearance and was unable to extrapolate to other rodent species^52^, our model incorporates additional mechanisms including endocytosis- and exocytosis-mediated tissue uptake/efflux, saturable host-response clearance, and size and shape-dependent parameters, and lymphatic drainage from all tissues. By estimating maximum macrophage uptake capacity from dose-ranging data and incorporating transport between vascular, interstitial, and cellular sub-compartments (informed by human cell culture studies), our model addresses prior limitations and improves predictive accuracy across tissues, phage families, and host species (**Fig. 9**).Phages can cross human epithelial barriers bidirectionally via endocytosis and exocytosis ^35^, and can undergo lysosomal degradation, as shown for *E. coli* phages K1F and PK1A2 in human epithelial and neuroblastoma cells ^36, 37^. Our PBPK model incorporates these processes and, in addition to MPS and lysosomal-mediated clearance, includes plasma clearance required to describe the observed data. This is consistent with reports of accelerated plasma clearance following repeated phage dosing without corresponding changes in liver or spleen phage titers (primary organs of MPS clearance), suggesting additional elimination pathways^41^. Such additional elimination pathways are unlikely to involve neutrophils as recent data indicates that neutrophils are not a major determinant of phage clearance, as shown by similar phage PK in neutropenic CD1 mice relative to BALB/c controls^53^. Together these findings highlight the complexity of phage elimination involving multiple mechanisms and underscore the need for continued work dissecting host determinants of phage clearance.

Clinical success with inhaled phage therapy including reduction in multidrug-resistant *Pseudomonas aeruginosa* and *Staphylococcus aureus* infection in a 7-year-old CF patient^54^ and favorable outcomes with nebulized phages including Muddy, BPsΔ, and/or ZoeJΔ support the clinical feasibility of inhaled delivery^20,28^. Our data provides mechanistic justification for this route: IT administration produced higher lung exposure with lower systemic exposure, a profile desirable for pulmonary NTM treatment. These findings are consistent with prior reports comparing IV and IT dosing of the 536-P1 phage^55^. Interestingly, our model indicates more rapid plasma clearance after IT administration (472 mL/h) than after IV (10.8 mL/h). To our knowledge, this is the first study to directly compare phage disposition across plasma, lung, liver, and spleen following both IV and IT administration.

External validation of our PBPK framework using diverse cross-species phage PK datasets identified size and morphology as critical PK determinants of phage disposition. Specifically, podophages, which are distinguished by their smaller size and short tails, exhibited higher plasma clearance compared to siphophages. Our successful model-based optimization provided further support of slower exocytosis rates for podophages, which was suggested previously by observations that T3 phage (podophage) accumulates intracellularly to a greater extent than T4 (myophage) and λ (siphophage) phages^42^.

Clinically used IV and nebulized phage doses (10⁶–10¹¹ PFU) remain largely empirically chosen. The ARLG Phage Taskforce noted that higher doses do not consistently yield superior outcomes relative to lower doses^56^, highlighting the need for exposure-guided dosing strategies. By enabling simulating human phage exposure in plasma and at the site of infection, such as the lung, our PBPK framework provides a foundation for model-informed dose and route selection. The model will also support sparse-sampling strategies for future PK studies that evaluate phage pharmacodynamics. Integration of this PBPK model with mechanistic phage–bacteria pharmacodynamic models will enable evaluation of “self-replicating” behavior of phage and refine dosing regimens that align phage amplification with bacterial kinetics.

In summary, this work delivers the first mechanistically resolved PBPK model of mycobacteriophage disposition; demonstrates route-dependent, saturable, and morphology-linked PK; and provides quantitative rationale supporting inhaled delivery for pulmonary Mabs infection. Our analysis also generated testable hypotheses including the existence of phage clearance mechanisms beyond MPS and lysosomal degradation and morphology-dependent differences in uptake and efflux that merit further studies. Collectively, these findings advance phage therapy from empirical derived regimens toward predictive, exposure-driven therapeutic design, accelerating clinical translation for multidrug-resistant NTM.

## Materials

### Bacterial strains, phage, and media

Lytic mycobacteriophage BPsΔ33HTH-HRM10 (BPsΔ), ZoeJΔ45 (ZoeJΔ), and Muddy were used in this study^20,27–29^. *Mycobacterium smegmatis* mc^2^155 was used as the bacterial host for phage amplification and quantification for the experiments. *M. smegmatis* was cultured on 7H10 agar (Difco), supplemented with oleic acid albumin dextrose complex (OADC) and 0.5 % glycerol^57^. Liquid cultures of *M. smegmatis* were grown in 7H9 (Difco) media containing 10 % albumin dextrose complex (ADC), 0. 5% glycerol, and 0.05 % Tween 80^57^.

### Mycobacteriophage Quantification and Production for PK Studies

Phage quantification and production of high titer phage stocks was performed following previously described methods with minor modifications^58,59^. Briefly, a 2-day culture of the bacterial host *M. smegmatis* mc^2^155 strain was prepared in 7H9 liquid medium at 37°C. Then, using the double agar overlay method, 200 μL of *M. smegmatis* bacteria and 100 μL of 10-fold serial dilutions of phage were mixed in 3.5 mL of molten top agar (Difco Agar Noble, BD Biosciences) and poured onto 7H10 agar plates. Plates were incubated at 37°C for ∼18 hours for BPsΔ and ZoeJΔ and for ∼48 hours for Muddy, which forms smaller plaques. Phages were quantified as plaque forming units (PFU) and the lower limit of quantification (LLOQ) is 1.3 log_10_ PFU/mL.

Phage dilutions generating a webbed appearance of phage plaques in the overlays were selected for preparing high titer stocks used in subsequent steps. Phages were harvested by flooding each plate with 5 mL of phage buffer (10 mM Tris, pH 7.5, 10 mM MgSO_4_, 68 mM NaCl, 1 mM CaCl_2_) and incubating at room temperature for 4 hours. The resulting lysate was filtered through a 0.2-μm syringe filter and precipitated overnight at 4℃ using 4% polyethylene glycol (PEG) 8000 (Fisher Scientific, US) in 0.5 M NaCl. The PEG pellet was collected by centrifugation for 30 min, 3,700 x *g* at 4℃ and resuspended in phage buffer by overnight incubation at 4℃. Following a second spin to remove any undissolved material, the phage-containing supernatant was transferred to an ultrafiltration centrifugation column with a 100 kDa molecular weight cutoff (Pierce Protein Concentrator, Thermo Fisher Scientific), washed with 40 mL of phage buffer,^60^ and concentrated to a smaller volume by centrifugation at 3,700 x *g* at 4°C. The concentrated phage was recovered from the upper reservoir of the concentrator, filtered through a 0.22-µm filter, and stored at 4° C. Prior to use in mouse PK experiments, the phage buffer was exchanged with sterile saline (Grifols, S.A. IV injection solution 0.9% Sodium chloride; Laboratories Grifols S.A.) by centrifugation with 100 kDa ultrafiltration centrifugation columns.

### In Vivo PK Studies

To evaluate the distribution and resulting exposure of mycobacteriophages BPsΔ, ZoeJΔ, and Muddy, PK studies were conducted in uninfected mice following intravenous (IV) or intratracheal (IT) administration. Female C3HeB/FeJ (6–8 weeks old, 22–25 g) were sourced from the Jackson laboratory (Bar Harbor, ME, USA). Female β-ENaC mice (C57Bl/6N *Scnn1b*-Tg mice^61^ (7-12 -weeks old) were bred at the Marsico Lung Institute Animal Models Core (now part of the UNC Pulmonary and Inhalation Research Core) and genotyped for Scnn1b-Tg expression by Transnetyx (CB6608 line). Mice were housed in individually ventilated micro-isolator cages within a specific pathogen-free facility at the University of North Carolina at Chapel Hill on a 12-hour light/dark cycle. Mice were provided with standard chow diet and water ad libitum. All procedures were approved by the Institutional Animal Care and Use Committees (IACUC) of both the University of North Carolina and the University of Southern California and conducted in accordance with the Animal Welfare and the National Institutes of Health (NIH) guidelines for the care and use of animals in biomedical research.

#### Dexamethasone Treatment

Select groups of mice from both strains (C3HeB/FeJ and β-ENaC) received subcutaneous injections of 5 mg/kg dexamethasone (DEX; 2 mg/mL; VetTek, Blue Springs, MO, USA.) beginning 7 days prior to phage administration and continuing throughout the duration of the study. Dexamethasone treatment was implemented to enable PK data collection under relevant immunosuppressive conditions that are used to establish stable Mabs infection in mice, as previously described^30^.

#### Intravenous (IV) and Intratracheal (IT) Dosing

Throughout this study, the high dose was 4 × 10^12^ PFU/kg, medium was 4 × 10^10^ PFU/kg, low was 4 × 10^7^ PFU/kg. Both IV and IT administration routes were evaluated for all phages (BPsΔ, ZoeJΔ, and Muddy) in C3HeB/FeJ mice. However, in β-ENaC, only BPsΔ was administered via the IV route. For IV administration, phages were injected via the tail vein (0.1 mL/mouse of concentrated phage in saline). For IT dosing, mice were first anesthetized with ketamine (100 mg/mL, Covetrus North America, OH, USA) at 100 mg/kg and dexmedetomidine (0.5 mg/mL, Dechra Veterinary Products, KS, USA) at 0.5 mg/kg delivered intraperitoneally (25 µL of each for a 25 g mouse). The trachea was visualized using an otoscope, and a silicon tube connected to a 500 µL syringe used to deliver 0.1 mL/mouse of concentrated phage in saline directly into the trachea under visual guidance. Following IT installation of phage, mice received 1 mg/kg of antisedan (atipamezole hydrochloride; Modern Veterinary Therapeutics, Florida, USA) to reverse the effect of sedatives.

#### Sampling and Phage Quantification

At predefined time points, groups of mice (n=3) were humanely euthanized via 100% carbon dioxide inhalation (30-70% CO_2_ displacement rate in the euthanasia chamber) followed by cardiac puncture. Heparinized blood (plasma) and tissues including lung, liver, and spleen were collected. Inguinal lymph nodes were also harvested in selected experiments. Tissues were placed in tubes containing phage buffer and homogenized using either a Polytron homogenizer (10,000 rpm; 15–30 seconds) or a MiniMix homogenizer (Interscience; 2 minutes). Homogenates were centrifuged, and the supernatant was transferred to vials for plaque-forming unit (PFU) quantification. All samples were kept on ice and processed within 1 hour of collection. Phage enumeration was performed using the double agar overlay method with serially diluted plasma and tissue homogenates as described above. PFU/mL (blood) or PFU/g (tissue) was calculated accordingly.

#### Phage PK Comparisons

PK studies were conducted for three distinct phages, BPsΔ, ZoeJΔ, and Muddy, administered at high and low doses via IV or IT routes in DEX-treated C3HeB/FeJ mice to assess differences in distribution and clearance.

#### Perfusion Study

A perfusion study was conducted in DEX treated C3HeB/FeJ mice (n=6) following high dose (4 × 10¹² PFU/kg) IV administration of BPsΔ phage to determine the influence of residual blood on phage quantification of tissue samples. At 4 hours post-phage administration, mice were divided into two groups: perfused and non-perfused. For the non-perfused, euthanasia was performed as described above and immediately followed by cardiac puncture and plasma and tissue collection. In the perfused group, following cardiac puncture, 10 mL of 1x PBS was slowly injected into the right ventricle until the lungs turned pale, indicating successful perfusion. Tissues from the two groups were homogenized and analyzed for phage concentration using the double agar overlay method as previously described.

Following lung perfusion, no significant difference was observed in the phage counts in plasma, lung, liver, and spleen (**Supplementary Fig. 3**), hence all the PK studies were performed in non-perfused mice.

### Non-compartmental analysis

In vivo PK data was analyzed using non-compartmental analysis (NCA) with PKanalix (version 2024R1, Lixoft SAS, France) software. The area under the concentration-time curve (AUC) was calculated using the linear trapezoidal method.

### Physiologically Based Pharmacokinetic (PBPK) Model Development

The biodistribution and elimination of BPsΔ in DEX-treated C3HFeBJ mice was characterized using a PBPK framework that includes i) circulation of plasma and lymph, ii) phage distribution to each organ, iii) interaction of phage with the host immune response, and iv) role of lymphatic system in the distribution of phage from tissues to the bloodstream.

Time-course PK data following BPsΔ phage administration via IV or IT were simultaneously modeled using a naïve pooled approach with the maximum likelihood estimation in ADAPT5 (Biomedical Simulations Resource, Los Angeles, CA, US). Data visualization and model simulations were performed in R (v4.5.1). Model performance was evaluated based on visual inspection of model fits and precision of parameter estimates.

### Model-Based Simulations of In vivo PK for ZoeJΔ and Muddy

We conducted simulations in 1,000 virtual mice following administration of a high dose of 4 × 10¹² PFU/kg of ZoeJ**Δ** or Muddy via IV or IT routes to assess the PBPK model’s ability to predict the PK of mycobacteriophages other than BPsΔ used to develop the model. Simulations were performed with 30% variability across parameters and a residual unexplained variability of 0.3 log_10_ PFU/mL.

### Interspecies Scaling of PBPK Model

To further validate the PBPK model, PK data from different phage families were extracted from published literature (**Supplementary Table 4**). Phage concentration–time profiles in plasma and/or tissues (lung, liver, and spleen) were digitized using WebPlotDigitizer (version 4.6). The dataset included phage concentration–time for mice, rats, and monkeys following IV administration, as well as data from mice following IT administration. Physiological parameters for mice, rats, and monkeys are provided in **Supplementary Table 2**, and detailed information regarding phage family, size, dose, and study design is summarized in **Supplementary Table 4**. Model parameters were either final model estimates or optimized to describe the observed data. Plasma clearance was scaled allometrically across species. Simulations were performed for 1,000 virtual subjects per species, assuming 30% parameter variability and residual unexplained variability ranging 0.3–1 log_10_ PFU/mL, depending on each study’s lower limit of quantification (LLOQ/2).

To assess the ability to scale phage dosing to humans, plasma concentration-time data were digitized for the *Escherichia coli* phage cocktail LBP-EC01 administered to patients with uncomplicated urinary tract infections via the IV route (**Supplementary Table 4)**^40^.

## Data availability

All data that support the findings of this study are provided in the article and the Supplementary material files.

## Acknowledgments

Procurement of *Scnn1b*-Tg mice by the Marsico Lung Institute Animal Models Core (now part of the UNC Pulmonary and Inhalation Research Core) was supported by Cystic Fibrosis Foundation Research Development Project BOUCHE19R0, National Institutes of Health grants NHLBI P01HL164320, and NIDDK P30 DK065988 to R.C.B, and NHLBI R01 HL150541 to A.L-B. This work was supported by NIAID R01 AI176834 to GRR, MB, SR and Cystic Fibrosis Foundation grant BRAUNS21P0 to MB and GRR. A.A.S. acknowledges support from a Kirschstein National Research Service Award (T32AI007151). This work was also supported by grants from the National Institutes of Health (GM116884), Howard Hughes Medical Institute (GT12053), Cystic Fibrosis Foundation (HATFUL19GO), and Emily’s Entourage (AWD00006613) to GFH.

## Author’s contribution

G.G.R. and M.B. conceived the study and secured funding. R.S., H.T., A.A.S., M.B., and G.G.R. designed the experiments. R.S., S.Y., D.S., C.C., H.T., A.A.S., and P.N. performed the in vivo studies. R.S., R.M., S.M., and G.G.R. analyzed the data and interpreted the results. G.G.R., R.S., R.M., and M.B. wrote the manuscript. A.A.S., S.E.M., A.J.M., and G.F.H. provided critical revisions and editorial input. All authors reviewed and approved the final manuscript.

